# RDEmbed: A Sentence-Embedding Framework for Rare Disease Phenotype Representation and Phenotypic Similarity Analysis

**DOI:** 10.64898/2026.10.16.759399

**Authors:** Xinghua Wang, Jinlian Wang, Zijian Zhang, Hongfang Liu, Arif Harmanci

**Affiliations:** Department of Health Data Science and Artificial Intelligence, D. Bradley McWilliams School of Biomedical Informatics, The University of Texas Health Science Center Houston, Houston, TX, 77030, USA; Department of Quantitative and Systems Health Sciences, The University of Texas at Austin Dell Medical School, Austin, TX, 78712, USA; Section of Epidemiology and Population Sciences, Department of Medicine, Baylor College of Medicine, Houston, TX, 77030, USA

**Keywords:** Rare disease, Phenotyping, Sentence embedding, Phenotypical similarity, Human Phenotype Ontology, Transformer model

## Abstract

Accurate phenotypic characterization of patients with rare diseases is essential for improving diagnosis, patient matchmaking, and gene discovery. Conventional ontology-driven approaches based on the Human Phenotype Ontology (HPO) enable structured phenotype comparison but often limited by the incomplete annotations and reduced ability to capture the contextual information contained in free-text clinical narratives.

We present RDEmbed, a sentence embedding framework that learns patient phenotype representations directly from rare disease case reports for similarity analysis. RDEmbed encodes narrative case reports into high-dimensional vectors to quantify phenotypic similarity. Unlike ontology-based approaches that rely on manually curated HPO annotations, RDEmbed encodes narrative case reports into dense semantic embeddings that capture complex phenotypic relationships from unstructured text. We further introduced retrieval-augmented embedding refinement (RAER), which enhances embedding representations by incorporating external domain knowledge at inference time without task-specific fine-tuning.

RDEmbed was evaluated against HPO-based similarity methods (OWLSim, simGIC) using simulated datasets, published rare disease case reports and a curated case study dataset comprising 156 patient abstracts presenting six rare disease groups. Across the real world datasets and curated cohort. RDEmbed consistently outperformed ontology-based approaches, improving AUROC and AUPRC by more than 5%. Embedding-based representations also produced more distinct disease-specific clustering in t-SNE visualizations and identified twice as many true positive rare disease pairs as ontology-based methods (8 versus 4). Incorporating RAER further improved precision while maintaining high sensitivity compared with zero-shot embeddings.

These results demonstrate that sentence embeddings provide an effective alternative to ontology-based phenotype representations by learning directly from free-text clinical narratives. RDEmbed enables more accurate phenotypic similarity analysis while reducing dependence on manually curated HPO annotations and offers a scalable framework for rare disease diagnosis, patient matchmaking, and gene discovery. RDEmbed is publicly available at https://github.com/xhwang2017/RDEmbed.

## Introduction

More than 7,000 distinct rare diseases have been identified, each affecting fewer than one in 2,000 individuals globally. Collectively, these disorders impact an estimated 300 million people, the majority of whom have conditions with a genetic basis, accounting for roughly 80% of cases [1]. Although next-generation sequencing has improved variant discovery, only 30–40% of rare disease cases currently achieve molecular diagnosis. [2]– [4]. For many rare disorders, detailed clinical case reports remain among the most valuable resources for disseminating new disease presentations, diagnostic strategies, and management outcomes [5]. These reports often contain detailed narrative accounts of patient phenotypes, diagnostic findings, and therapeutic responses that, when systematically analyzed, can uncover shared clinical patterns across individuals [6]–[8]. Effective computational phenotyping of such text is therefore essential for identifying phenotypically similar cases, enabling patient matchmaking, and ultimately improving diagnostic accuracy in rare disease settings.

The Human Phenotype Ontology (HPO) has become the standard framework for representing phenotypic abnormalities in rare disease research [9], [10]. By organizing standardized clinical features into a hierarchical ontology, HPO enables computational comparison of patient phenotypes using semantic similarity metrics such as Resnik, Jiang–Conrath, Wang, simGIC, and OWLSim [11]–[15]. These methods underpin widely used tools including Exomiser, PhenoDigm, and PhenomeCentral for gene prioritization, disease diagnosis, and patient matchmaking [16]–[18]. Despite their success, HPO-based phenotyping approaches face notable constraints. Their performance relies on accurate extraction of HPO terms from unstructured text, which remains error-prone even with advanced natural language processing tools [19], [20]. Moreover, these ontology-derived similarity scores are limited by the completeness of annotations, the quality of evidence codes, and the inability of information content–based metrics to fully capture the nuanced semantic context present in narrative case descriptions [21], [22]. Consequently, clinically meaningful contextual information contained within narrative case descriptions, including disease progression, phenotype interactions, diagnostic reasoning, and atypical manifestations,may not be fully captured. These limitations can reduce the sensitivity of patient similarity analyses and hinder the discovery of previously unrecognized disease relationships.

Recent advances in large language models (LLMs) and biomedical natural language processing have substantially expanded the computational phenotyping landscape beyond traditional ontology-driven approaches. Early domain-adapted models such as BioBERT [23] and ClinicalBERT [24] demonstrated significant improvements in biomedical named entity recognition, relation extraction, and clinical concept normalization. Building on this foundation, sentence-level transformers [25] generate embeddings for each sentence or short text segment, enabling fine-grained semantic comparisons. Unlike ontology-based methods that rely on explicit phenotype annotations, these embeddings allow clinical narratives, including case reports, to be projected into a continuous semantic vector space, where cosine similarity can quantify phenotypic similarity without requiring explicit HPO annotations [26].

The diversity of contemporary computational phenotyping approaches is summarized in Table 1. Existing methods can be broadly categorized into ontology-grounded phenotype extraction systems and embedding-based semantic representation models. Ontology-centered approaches, including PhenoBERT, PhenoHPO, FastHPOCR, and RAG-HPO, focus on extracting and normalizing clinical concepts into standardized HPO terms [27]– [30]. These methods provide highly interpretable outputs and facilitate integration with established rare disease analysis pipelines. More recent LLM-based systems such as PhenoGPT and SemEHR+LLaMA3 extend phenotype extraction through generative reasoning and broader disease concept recognition [20], [31]. However, these approaches ultimately depend on explicit phenotype identification and ontology-defined representations, which may overlook latent semantic relationships contained within narrative clinical descriptions.

**Table 1.** Comparison of Computational Phenotyping Approaches for Rare Disease Research.

| Method | Model | Representation | Coverage | Interpretability | Task |
| --- | --- | --- | --- | --- | --- |
| PhenoBERT[32] | BERT + CNN | Explicit HPO terms | Limited to defined HPO | High | HPO term recognition |
| PhenoHPO[28] | Fine-tuned Llama-2 | Explicit HPO terms | Limited to defined HPO | High | Concept normalization |
| FastHPOCR[29] | Dictionary-based | Ontology-grounded symbolic HPO terms | HPO-defined phenotypes | High | Phenotype concept recognition from biomedical abstracts |
| RAG-HPO[30] | LLaMa-3.1 + RAG | Explicit HPO terms | Defined HPO + synonyms | High | Accurate HPO term assignment |
| PhenoGPT[20] | GPT-based LLM | HPO terms and expanded new phenotypes | Moderate (HPO+new phenotype) | Moderate | Phenotype detection & vocab expansion |
| SemEHR+LLaMA3[31] | Dictionary-based NLP + LLaMA3 | Rare disease mentions | High | High | Rare disease mentions identification |
| RDEmbed | All-MiniLM-L6-v2 | Dense vector | Very High<br>(latent info) | Low | Clustering,<br>patient<br>similarity<br>matching |

To address these limitations, we developed RDEmbed, an embedding-based framework for rare disease phenotyping and patient similarity analysis (Figure 1A). RDEmbed utilizes a pre-trained sentence-transformer model to encode clinical narratives directly into dense semantic vectors without requiring phenotype extraction, ontology mapping, or task-specific model fine-tuning. By representing case reports in a continuous semantic space, phenotypic similarity can be quantified through embedding-based similarity measures that preserve contextual information contained within narrative clinical descriptions.

**Figure 1.**
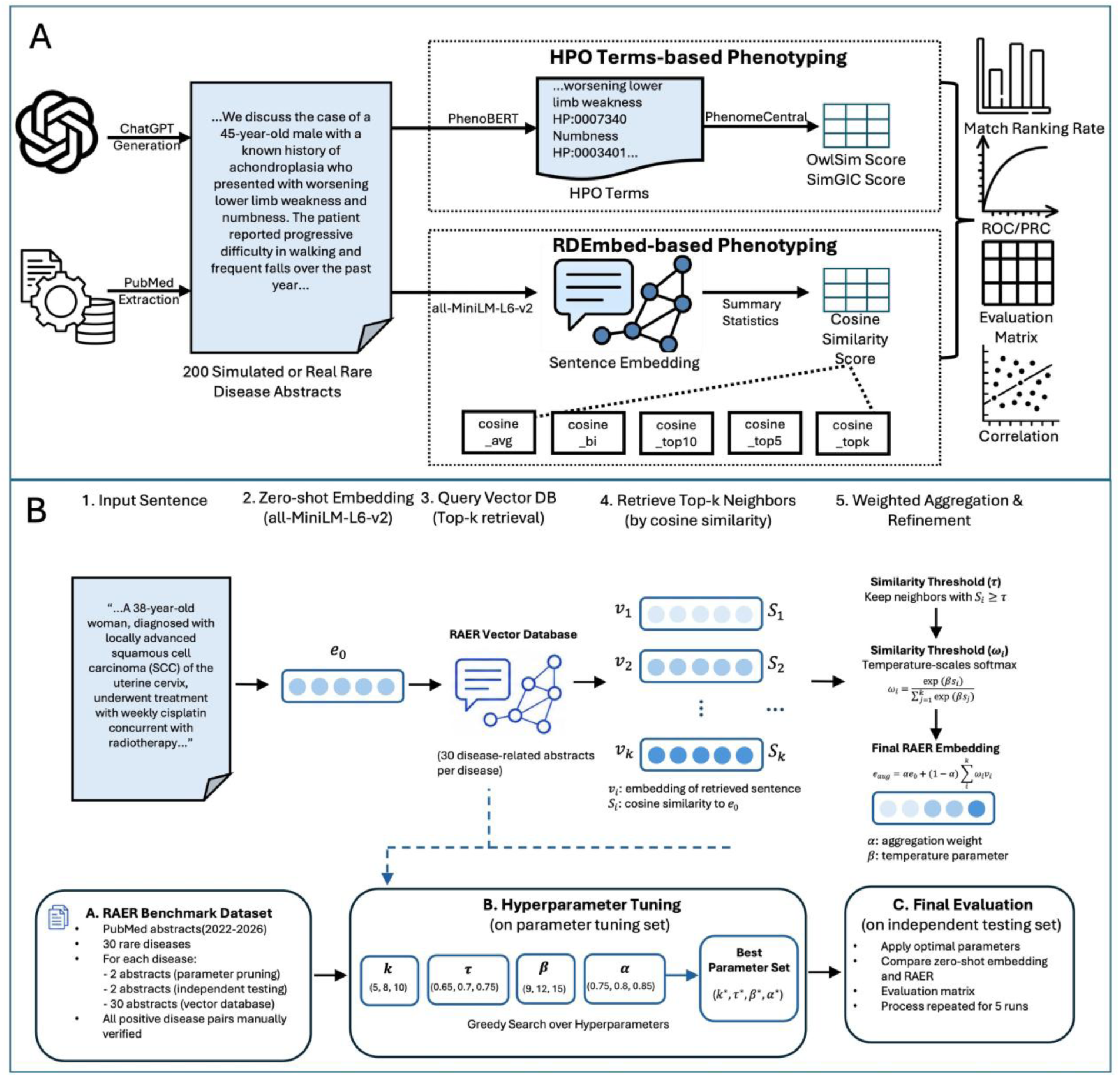
a. Overview of the Study Workflow. A simulated dataset comprising 200 PubMed-style case report abstracts was generated for 100 rare diseases from Orphanet, alongside 200 additional real PubMed abstracts. Phenotypic features were extracted using PhenoBERT, and HPO-based semantic similarity was computed with OWLSim and SimGIC via PhenomeCentral. For comparison, RDEmbed, a transformer-based framework utilizing sentence-embedding cosine similarity derived from *all-MiniLM-L6-v2*, was applied to the same dataset. Rare disease matching performance was evaluated by ranking true disease pairs and computing ROC and PR curves, as well as accuracy, precision, recall, and F1 scores. This design enables a direct and systematic comparison between embedding-based and HPO-based phenotyping approaches using both standardized simulated data and real-world case reports. Figure 2B**. Retrieval-Augmented Embedding Refinement (RAER) Pipeline.** Given an input rare disease abstract, a zero-shot embedding is first generated using the pretrained all-MiniLM-L6-v2 model. The embedding is then used to query a vector database containing disease-related abstracts, and the top-*k* most similar neighbors are retrieved based on cosine similarity. Retrieved neighbors are filtered using a similarity threshold (*τ*) and weighted according to their relevance to the query. The filtered neighbors are subsequently aggregated with the original embedding to generate a refined retrieval-augmented embedding. Hyperparameters, including the number of retrieved neighbors (*k*), similarity threshold (*τ*), temperature parameter (*β*), and aggregation weight (*α*), were optimized using a dedicated parameter optimization dataset. The final model was evaluated on independent testing datasets across five repeated runs to assess robustness and generalizability.

Although zero-shot embeddings capture broad semantic relationships, they may not fully leverage disease-specific knowledge contained within the biomedical literature. Many recent approaches seek to improve phenotypic analysis through model fine-tuning, in which pre-trained language models are further optimized using domain-specific datasets [20], [28], [33]. While fine-tuning can enhance task performance, it often requires substantial computational resources, specialized expertise, and extensive training data. Moreover, performance gains may be inconsistent across tasks and model architectures, particularly for large foundation models [34]–[38]. Given the rapid evolution of language models and the continual expansion of phenotype ontologies such as HPO, maintaining and updating fine-tuned models can be challenging, limiting their practicality for many rare disease researchers and clinicians.

Retrieval-augmented generation (RAG) has emerged as a scalable alternative by incorporating external knowledge at inference time rather than modifying model parameters [39]–[41]. By retrieving relevant information from vector databases, RAG systems can continuously leverage newly available biomedical knowledge while avoiding costly retraining. Recent studies, including RAG-HPO, have demonstrated that retrieval augmentation can improve phenotype extraction and HPO term assignment from clinical narratives [30]. Building on this concept, we developed Retrieval-Augmented Embedding Refinement (RAER) (Figure 1B), which retrieves semantically related rare disease documents and integrates their embeddings with the original query representation through similarity-weighted aggregation. Unlike retrieval-augmented phenotype extraction methods, RAER directly refines semantic embeddings, enriching disease-specific contextual information to improve patient similarity estimation while preserving the efficiency, flexibility, and generalizability of zero-shot inference.

Despite rapid progress in biomedical language models, sentence embedding methods remain relatively underexplored for rare disease phenotyping, and retrieval-augmented embedding refinement has not been systematically evaluated for patient similarity analysis. Therefore, this study aims to (1) evaluate the effectiveness of zero-shot sentence embeddings for rare disease phenotypic similarity analysis, (2) develop and assess a retrieval-augmented embedding refinement strategy for improving semantic patient representations, and (3) benchmark embedding-based approaches against established HPO-based methods across simulated and real-world rare disease datasets. By providing a comprehensive comparison of ontology-based and embedding-based phenotyping strategies, this work establishes a practical framework for semantic patient matching and advances computational approaches for rare disease diagnosis.

## Results

Figure 1 shows overview of our study design. We make extensive use of abstracts from simulated and real-world rare disease case reports as our main data resource for simulations and benchmarking. For each patient, the sentences are embedded (See Methods) and patient-level summarized, followed by comparison via pairwise cosine similarity between all cases. We finally compare HPO terms-based similarity metrics (top-k matching for each method) using comparable patient-level summarization of the terms, specifically, OWLSim and simGIC [14]. We compared embedding-derived cosine similarity measures with HPO-based similarity scores using a simulated dataset of PubMed-style abstracts, a real-world corpus of published case reports, and a curated corpus of rare disease case report abstracts. By evaluating ranking accuracy, clustering behavior, and threshold-based sensitivity, we compare the embedding-based approaches and HPO-driven methods.

### t - SNE Visualization

To evaluate disease-specific clustering, we visualized 100 pairs of simulated abstracts, each pair describing the same rare disease. Sentence embeddings derived with the all-MiniLM-L6-v2 model were projected into two dimensions using t-SNE (Fig. 2A). Most paired abstracts clustered closely together, reflecting strong semantic alignment captured by the embeddings. In contrast, t-SNE plots based on HPO-term phenotyping (Fig. 2B) showed weaker pairing and less distinct clustering.

**Figure 2.**
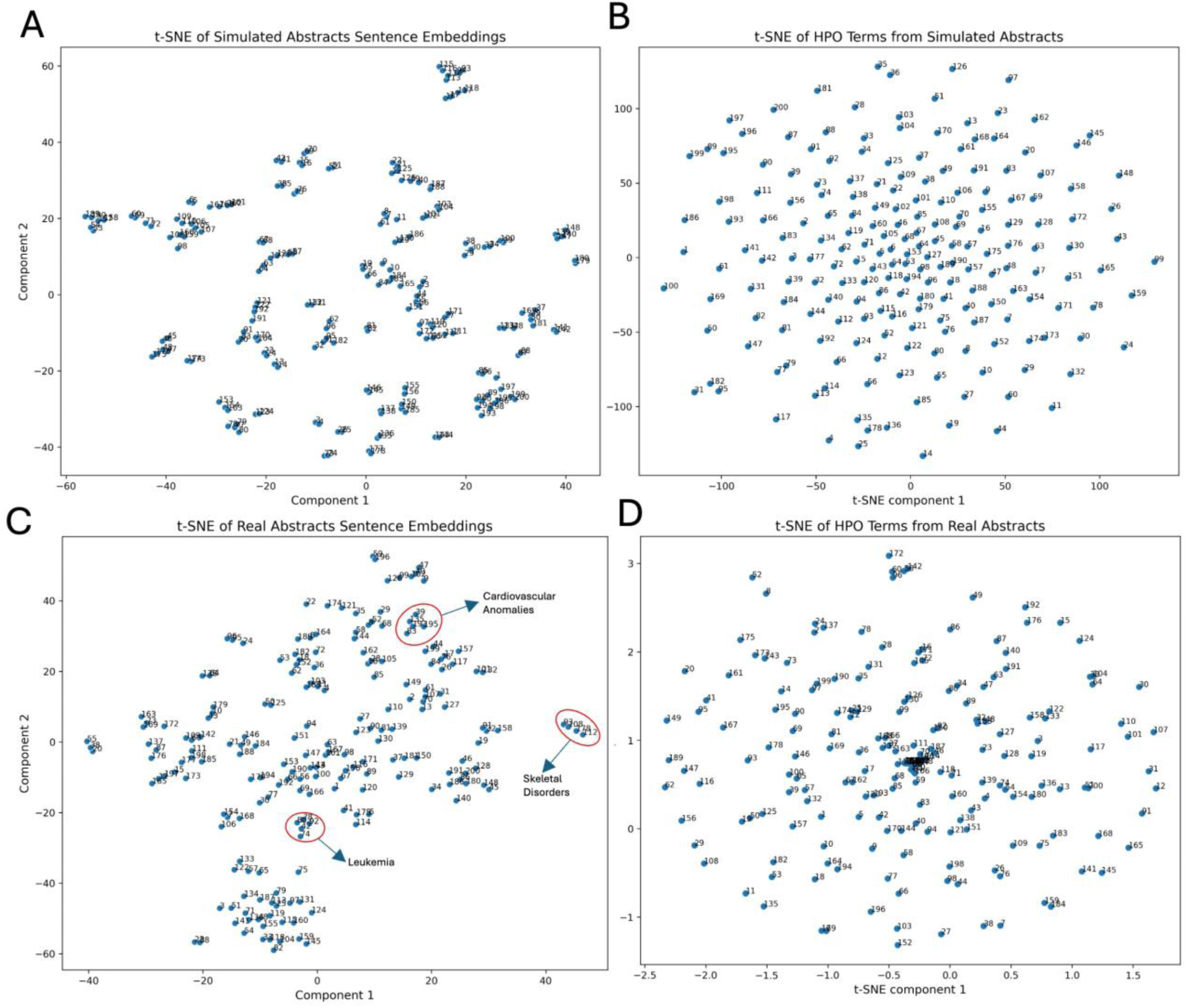
t-SNE visualization of rare-disease abstract clusters. **(A)** Sentence-level embeddings of 200 simulated abstracts generated with the all-MiniLM-L6-v2 model. **(B)** HPO term profiles derived from the same simulated abstracts. **(C)** Sentence-level embeddings of 200 real rare-disease abstracts generated with the all-MiniLM-L6-v2 model. **(D)** HPO term profiles derived from the corresponding real abstracts. In all panels, each point represents a single abstract or HPO profile, and numerals indicate sample indices. Clustering reflects similarities in semantic content or phenotypic annotation.

A parallel analysis of 200 real rare-disease case reports retrieved from PubMed yielded similar trends. t-SNE of sentence embeddings revealed multiple well-defined clusters of semantically related abstracts (Fig. 2C), whereas HPO-based projections were comparatively diffuse (Fig. 2D). Manual review confirmed that several tight clusters corresponded to shared clinical entities, including cardiovascular anomalies (abstracts 39, 83, 135, 193, 195), various leukemias (42, 74, 78, 80, 92), and skeletal disorders (93, 108, 112, 178). Collectively, these findings demonstrate that transformer-based sentence embeddings provide a more discriminative representation than HPO-based phenotyping, enabling clearer visualization of clinically coherent rare-disease clusters and more accurate identification of abstracts describing the same disease.

### Comparison of Similarity Scores for Simulated Abstracts

We evaluated the performance of each similarity measure on 100 pairs of simulated abstracts by computing the proportion of cases in which the correct rare disease match was ranked first (Top1) or appeared within the top five matches (Top5), as shown in Figure 3A (Supplementary Fig. S1, S2). Among HPO-based methods, OWLSim ranked the correct disease first in 42.5% of cases and within the top five in 70%, while simGIC exhibited higher accuracy in matching with 48% and 82.5%, respectively. Embedding-based approaches consistently outperformed these methods: Average cosine similarity (cosine_avg) ranked the true match first in 61% of pairs and within the top five in 87%. Adding bidirectional matching (cosine_bi) further increased these values to 87% and 98.5%. The highest-match similarity measures, cosine_top5, cosine_top10, and cosine_topk, demonstrated the best performance, correctly ranking the true disease first in 93% of cases and within the top five for nearly all pairs (99.5% for cosine_top5 and cosine_topk, 95.5% for cosine_top10). These results demonstrate that embedding-based similarity, particularly the highest-match measures, provides superior rare-disease identification compared with traditional HPO-based approaches.

**Figure 3.**
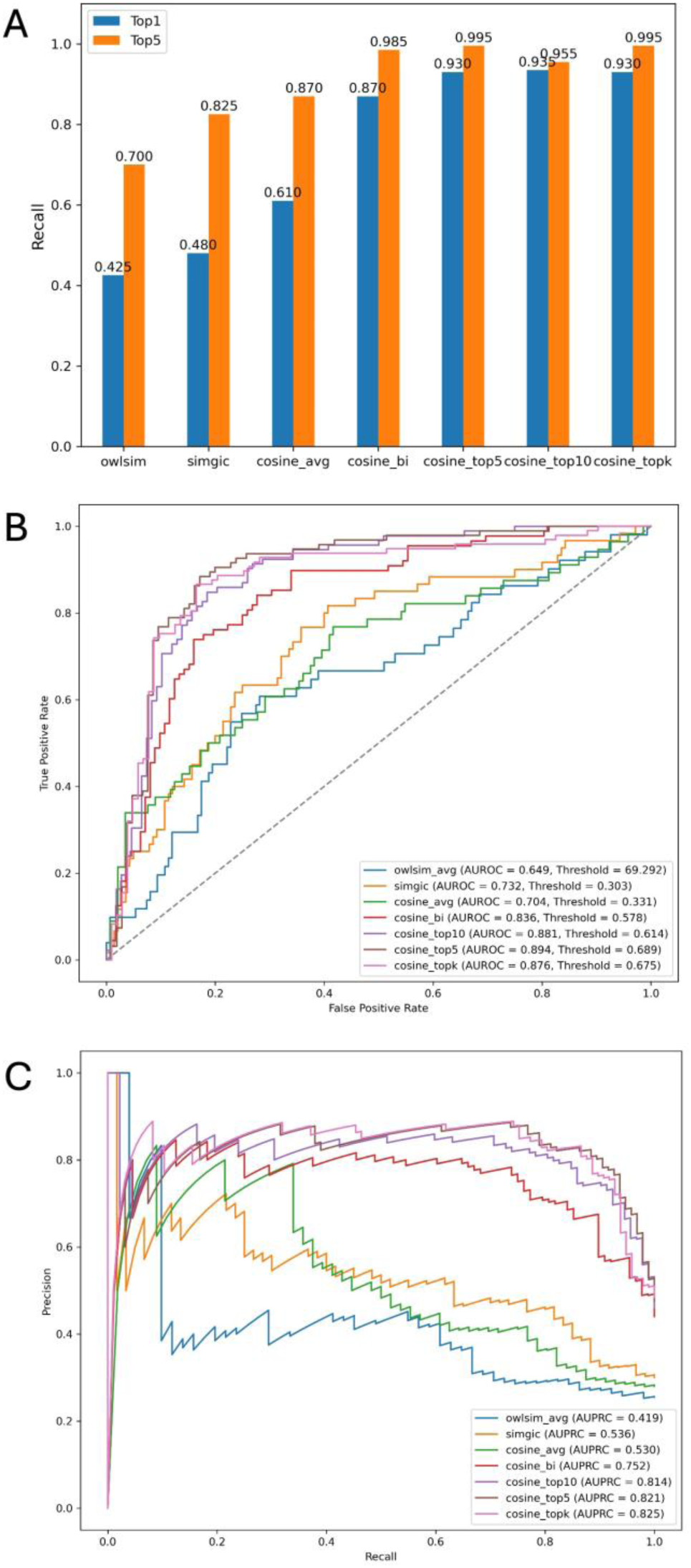
Performance comparison of seven similarity measures. **(A)** Fraction of cases in which the correct rare disease match was ranked first (Top1, blue) or within the top five (Top5, orange) most similar cases. Performance was evaluated on 100 pairs of simulated abstracts. **(B)** Receiver operating characteristic (ROC) curves for the seven similarity measures across the 100 simulated abstract pairs. **(C)** Precision–recall curves (PRC) for the seven similarity measures across the same dataset.

Seven similarity measures were evaluated on the simulated abstract dataset using accuracy, precision, recall, F1-score, AUROC, and AUPRC (Table 2). Among HPO-based approaches, OWLSim achieved an accuracy of 0.690 with moderate precision (0.425) and recall (0.608), while simGIC showed the highest precision (0.479), recall (0.767), and AUROC (0.732) within this group. Embedding-based methods exhibited concordance in terms of direction (Supplementary Fig. S3) and consistently outperformed HPO-based scores. The average cosine similarity (cosine_avg) had comparable AUROC (0.704) but lower accuracy (0.635). Incorporating bidirectional matching (cosine_bi) substantially improved performance, with accuracy 0.795, F1-score 0.760, AUROC 0.836, and AUPRC 0.752. The highest match embedding measures, cosine_top5 and cosine_topk, achieved the best results across all metrics. Cosine_top5 reached the highest accuracy (0.855), recall (0.884), F1-score (0.853), and AUROC (0.894), while cosine_topk obtained the highest precision (0.832) and AUPRC (0.825). Cosine_top10 also performed strongly, with accuracy 0.830 and AUROC 0.881. ROC and PRC for all measures are shown in Figure 3B and Figure 3C, respectively. These results indicate that embedding-based highest-match measures, particularly cosine_top5 and cosine_topk, provide the most accurate discrimination of true disease matches.

**Table 2.** Performance Comparison Across Similarity Scores for 100 Pairs of Simulated Abstracts.

| Scores | Accuracy | Precision | Recall | F1 | AUROC | AUPRC |
| --- | --- | --- | --- | --- | --- | --- |
| owlsim | 0.690 | 0.425 | 0.608 | 0.500 | 0.649 | 0.419 |
| simgic | 0.680 | 0.479 | 0.767 | 0.590 | 0.732 | 0.536 |
| cosine_avg | 0.635 | 0.417 | 0.768 | 0.541 | 0.704 | 0.530 |
| cosine_bi | 0.795 | 0.783 | 0.739 | 0.760 | 0.836 | 0.752 |
| cosine_top10 | 0.830 | 0.796 | 0.848 | 0.821 | 0.881 | 0.814 |
| cosine_top5 | **0.855** | 0.824 | **0.884** | **0.853** | **0.894** | 0.821 |
| cosine_topk | 0.850 | **0.832** | 0.866 | 0.848 | 0.876 | **0.825** |

### Threshold-Based Evaluation

To make hard calls on patient matching (“matching”/“non-matching” decision), we selected optimal distance decision thresholds derived from the ROC curves by maximizing Youden’s J statistic, selecting the point on each ROC curve that maximizes *(sensitivity + specificity − 1)* to achieve the best balance between true positive and true negative rates [27]. Based on this criteria, the distance cutoffs were selected as 0.693 for OWLSim, 0.385 for simGIC, 0.324 for cosine_avg, 0.587 for cosine_bi, 0.600 for cosine_top5, 0.696 for cosine_top10, and 0.672 for cosine_topk. We again observed significant correlation in HPO and embedding based scores (Supplementary Fig. S4) with similar distributions of scores for all methods (Supplementary Figure S5). Since simGIC and cosine_top5 were the top-performing scores within their respective HPO-based and embedding-based methods, we used these thresholds to compare their precision in identifying true matches across both datasets and to examine the correlation between their scores (Figure 4).

**Figure 4.**
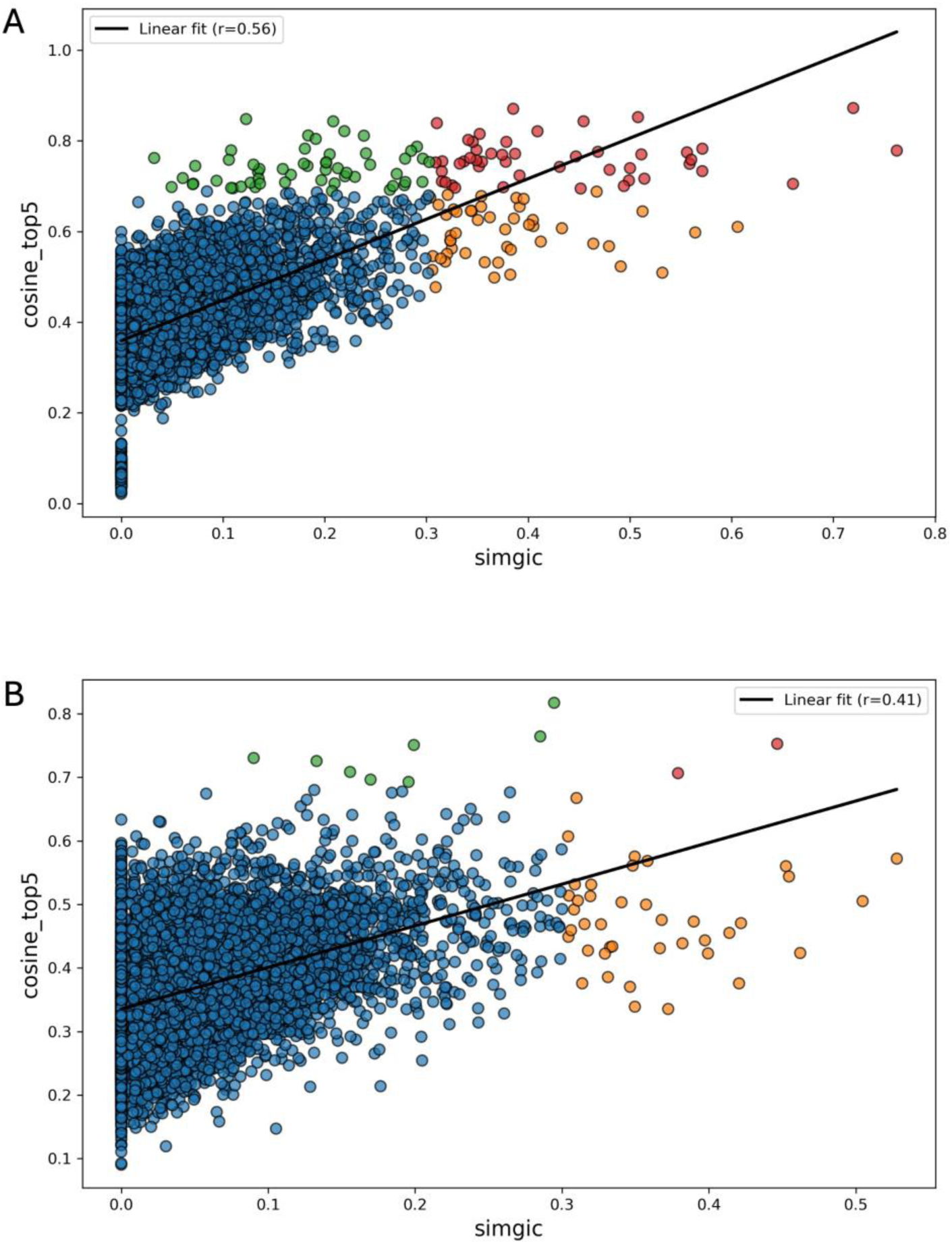
Correlation between SimGIC and Top5 Cosine similarity. **(A)** 100 pairs of simulated abstracts. **(B)** 200 real abstracts. Point colors indicate classification based on both similarity measures: **Blue:** Negative pairs identified by both SimGIC and Top5 Cosine similarity (simgic < 0.303 and cosine_top5 < 0.696). **Red:** Positive pairs identified by both SimGIC and Top5 Cosine similarity (simgic ≥ 0.303 and cosine_top5 ≥ 0.696). **Green:** Positive pairs identified by Top5 Cosine similarity (cosine_top5 ≥ 0.696) but negative by SimGIC (simgic < 0.303). **Orange:** Negative pairs identified by Top5 Cosine similarity (cosine_top5 < 0.696) but positive by SimGIC (simgic ≥ 0.303).

For both simulated and real datasets, cosine_top5 similarity consistently identified a greater number of true positive pairs compared to simGIC, indicating superior precision. In the simulated dataset (Fig. 4A), 47 pairs were classified as positive by both measures (red points), of which 39 were confirmed as true positives upon manual review. Fifty-five pairs were identified as positive by cosine_top5 but negative by simGIC (green points), with 45 validated as true positives. Conversely, 48 pairs were identified as positive by simGIC but negative by cosine_top5 (orange points), of which only 6 represented true positives. Based on these results, Top5 Cosine achieved a precision of 0.82, whereas simGIC had a precision of 0.47 in the simulated dataset.

In the real dataset (Figure 4B), two abstract pairs were classified as positive by both measures, both of which were true positives. Eight pairs were identified as positive by cosine_top5 but missed by simGIC, six of which were true positives. In contrast, 40 pairs were flagged as positive by simGIC but missed by cosine_top5, with only two confirmed as true positives. Table 3 lists these confirmed rare-disease abstract pairs, showing their HPO-based (simGIC) and embedding-based (cosine_top5) similarity scores and highlighting the greater precision of cosine_top5 in detecting true matches.

**Table 3.** True Positive Abstract Pairs Identified by simGIC and Cosine_top5.

| Abstract A | Abstract B | simgic | cosine_top5 | Rare Disease Involved | Identified by |
| --- | --- | --- | --- | --- | --- |
| P008 | P052 | 0.447 | 0.715 | Infective endocarditis with patent ductus arteriosus and pulmonary-artery vegetations | simgic & cosine_top5 |
| P108 | P178 | 0.379 | 0.676 | Femoral-head osteonecrosis (avascular necrosis) with fractures | simgic & cosine_top5 |
| P095 | P096 | 0.295 | 0.817 | Adrenal medulla tumor secreting excess catecholamines | cosine_top5 |
| P042 | P092 | 0.285 | 0.764 | Chronic lymphocytic leukemia | cosine_top5 |
| P059 | P196 | 0.199 | 0.751 | Spontaneous coronary-artery dissection (SCAD) causing acute coronary syndrome | cosine_top5 |
| P042 | P074 | 0.090 | 0.730 | Hemophagocytic lymphohistiocytosis (HLH) with atypical presentation | cosine_top5 |
| P093 | P108 | 0.133 | 0.726 | Missed or delayed femoral-neck fractures | cosine_top5 |
| P042 | P078 | 0.156 | 0.708 | Chronic lymphocytic leukemia with life-threatening complications | cosine_top5 |
| P036 | P182 | 0.196 | 0.693 | Transthyretin amyloidosis (ATTR) cardiomyopathy | cosine_top5 |
| P058 | P144 | 0.455 | 0.537 | Acute cardiac events from undiagnosed coronary artery disease during surgery | simgic |
| P010 | P053 | 0.452 | 0.525 | Peripartum cardiomyopathy (PPCM), including a case revealing undiagnosed sickle-cell disease | simgic |

These results indicate that HPO-based scoring exhibits lower sensitivity and precision relative to embedding-based highest-match measures. The correlation between simGIC and cosine_top5 scores was moderate (r=0.56 for simulated and 0.41 for real abstracts), suggesting that while the two methods capture overlapping phenotypic information, embedding-based measures provide complementary discriminative power, particularly in identifying true disease matches across diverse abstract datasets.

### Case Study for Curated Dataset

To evaluate the ability of the embedding-based approach to capture disease-specific phenotypic patterns, we applied t-SNE to visualize sentence-level embeddings and HPO matrix from 156 rare disease abstracts across six groups: Cystinuria (7 abstracts), Dent disease (11 abstracts), Primary Hyperoxaluria (PH; 13 abstracts), Idiopathic Pulmonary Fibrosis (IPF; 19 abstracts), Mastocytosis (47 abstracts), and Hypereosinophilic Syndrome (HES; 59 abstracts) (Supplementary Fig. S6). As shown in Figure 5A, the six groups generally formed cohesive clusters that were largely separable from each other, indicating that the embedding model effectively captured key phenotypic distinctions. Some overlap between the IPF and HES clusters, however, suggests the presence of shared or less distinct phenotypic features in the textual descriptions. By comparison, HPO-based profiles (Fig. 5B) exhibited weaker separation, highlighting the superior discriminative power of embedding-derived representations for rare disease phenotyping.

**Figure 5.**
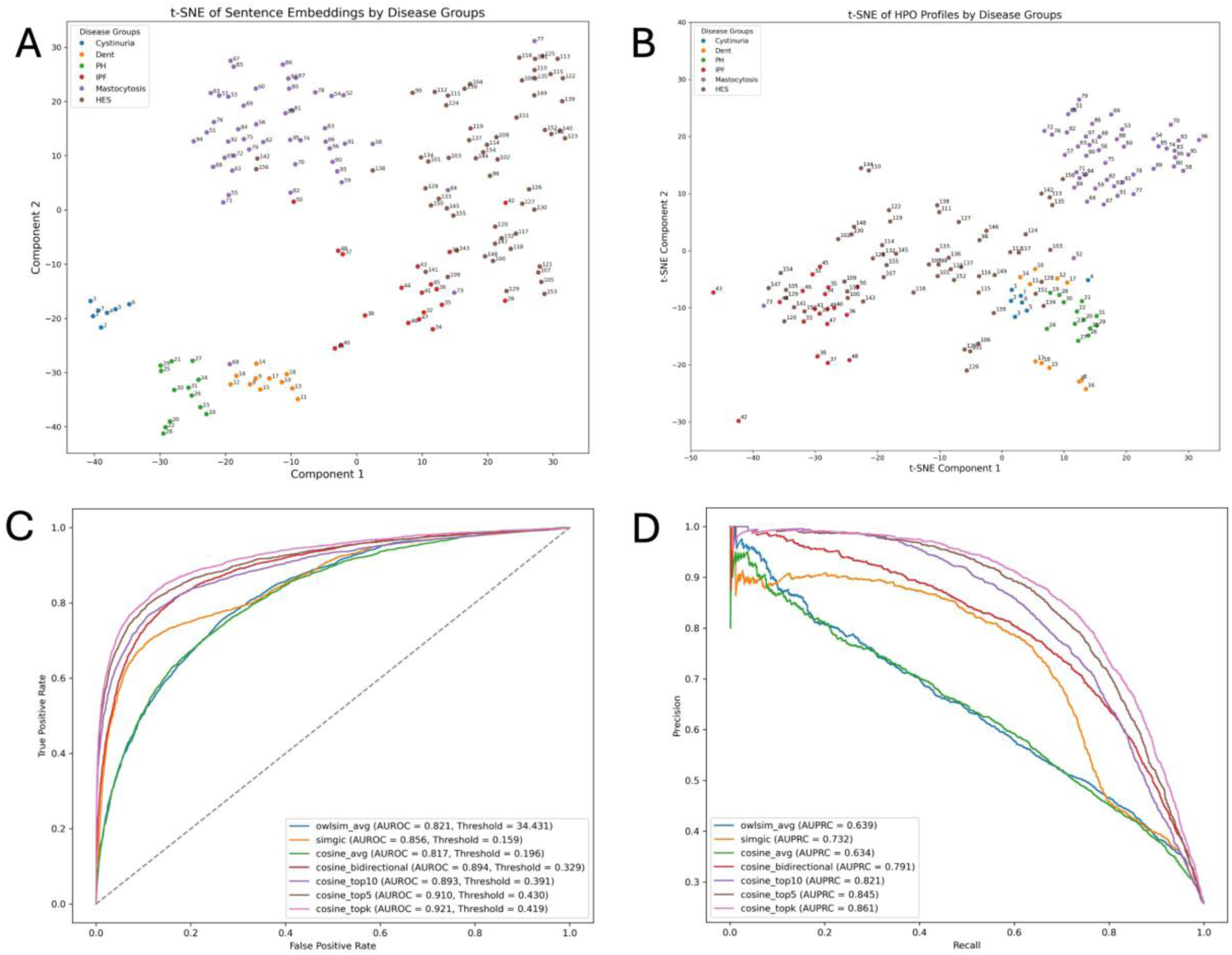
t-SNE visualization and performance comparison of seven similarity metrics on six curated groups of rare disease abstracts. **(A)** t-SNE visualization of sentence-level embeddings generated using the all-MiniLM-L6-v2 model. (B) t-SNE visualization of HPO term profiles derived from the same set of abstracts. (C) Receiver operating characteristic (ROC) curves for the seven similarity measures. (D) Precision– recall (PR) curves for the seven similarity measures.

Methods showed concordance in the phenotypic similarity trends in correlation analysis (Supplementary Fig. S7). Quantitative evaluation confirmed the superior performance of embedding-based similarity metrics compared with traditional HPO-based methods (Table 4). Among the HPO-based measures, SimGIC outperformed OWLSim in both accuracy (0.839 vs. 0.739) and F1-score (0.692 vs. 0.599), with corresponding AUROC/AUPRC values of 0.856/0.732 for SimGIC and 0.821/0.639 for OWLSim. Except for cosine_avg, all other cosine embedding–based metrics surpassed the HPO-driven methods across all evaluation criteria. Specifically, cosine_top5 achieved the highest accuracy (0.868) and precision (0.721), while cosine_topk achieved the highest recall (0.825) and F1-score (0.761). These metrics also demonstrated superior discriminative ability, with AUROC of 0.921 and AUPRC of 0.861. Overall, these results indicate that embedding-based phenotyping provides more consistent and discriminative similarity estimation between rare disease phenotypes than HPO-based methods.

**Table 4.** Performance Comparison Across Similarity Scores for Six Curated Groups of Rare Disease Abstracts.

| Scores | Accuracy | Precision | Recall | F1 | AUROC | AUPRC |
| --- | --- | --- | --- | --- | --- | --- |
| owlsim | 0.739 | 0.496 | 0.757 | 0.599 | 0.821 | 0.639 |
| simgic | 0.839 | 0.681 | 0.703 | 0.692 | 0.856 | 0.732 |
| cosine_avg | 0.752 | 0.513 | 0.711 | 0.596 | 0.817 | 0.634 |
| cosine_bi | 0.828 | 0.628 | 0.812 | 0.708 | 0.894 | 0.791 |
| cosine_top10 | 0.858 | 0.705 | 0.769 | 0.736 | 0.893 | 0.821 |
| cosine_top5 | **0.868** | **0.721** | 0.794 | 0.756 | 0.910 | 0.845 |
| cosine_topk | 0.866 | 0.706 | **0.825** | **0.761** | **0.921** | **0.861** |

### Retrieval-Augmented Embedding Refinement Performance

We evaluate a retrieval-augmented embedding refinement (RAER) framework for rare disease matching without task-specific fine-tuning. In biomedical informatics, fine-tuning large language or embedding models typically requires substantial computational resources, large annotated datasets, and considerable training time, yet may provide only limited or unstable performance gains [35], [37]. RAER addresses these limitations by incorporating external knowledge via retrieval at inference time to refine embedding representations without model retraining.

A benchmark dataset of 30 rare diseases was constructed from PubMed abstracts published between 2022 and 2026. For each disease, two abstracts were used for hyperparameter tuning and two for independent testing. Additionally, 30 disease-related abstracts per disease were stored in a vector database to support retrieval. Hyperparameters, including the number of retrieved neighbors k ∈ {5, 8, 10}, similarity threshold τ ∈ {0.65, 0.70, 0.75}, temperature β ∈ {9, 12, 15}, and aggregation weight α ∈ {0.75, 0.80, 0.85}, were optimized using greedy search on the tuning set. The best configuration was evaluated on the held-out test set. RAER was compared against a zero-shot baseline using the all-MiniLM-L6-v2 embedding model, with performance assessed by accuracy, precision, recall, F1-score, AUROC, and AUPRC.

Table 5 summarizes the results of the parameter pruning stage. Model configurations were selected based on AUPRC and precision, and classification thresholds were determined on the tuning set using Youden’s J statistic. The optimal thresholds (0.512 for the zero-shot baseline and 0.555 for RAER) were subsequently fixed and applied to the independent test set. Across all evaluated configurations, RAER consistently outperformed the zero-shot baseline. The top-performing configurations achieved a precision of 0.882 and an AUPRC of 0.993, compared with 0.714 and 0.971 for the zero-shot baseline, respectively. Recall remained at 1.0 across all configurations. Performance was stable across variations in the number of retrieved neighbors (k) and temperature parameter (β), suggesting that RAER is robust to moderate hyperparameter changes when the similarity threshold (τ) and aggregation weight (α) are fixed. Results for all evaluated configurations are provided in the Supplementary Table 1.

**Table 5.** Parameter Tuning Results for RAER Compared with the Zero-Shot Baseline.

| Model/Configuration | Accuracy | Precision | Recall | F1 | AUROC | AUPRC |
| --- | --- | --- | --- | --- | --- | --- |
| Zero-shot | 0.993 | 0.714 | 1.0 | 0.833 | 0.999 | 0.971 |
| k8_τ0.70_β9_α0.75 | **0.998** | **0.882** | 1.0 | **0.938** | **1.0** | **0.993** |
| k8_τ0.70_β12_α0.75 | 0.998 | 0.882 | 1.0 | 0.938 | 1.0 | 0.993 |
| k8_τ0.70_β15_α0.75 | 0.998 | 0.882 | 1.0 | 0.938 | 1.0 | 0.991 |
| k10_τ0.70_β9_α0.75 | 0.998 | 0.882 | 1.0 | 0.938 | 1.0 | 0.993 |
| k10_τ0.70_β12_α0.75 | 0.998 | 0.882 | 1.0 | 0.938 | 1.0 | 0.993 |
| k10_τ0.70_β15_α0.75 | 0.998 | 0.882 | 1.0 | 0.938 | 1.0 | 0.991 |

The selected configurations were subsequently evaluated on an independent test set using the same parameter settings and thresholds. As shown in Table 6, RAER maintained superior performance relative to the zero-shot baseline, improving precision from 0.674 to 0.778 and F1-score from 0.795 to 0.848. Although recall decreased modestly from 0.967 to 0.933, the gain in precision indicates a reduction in false-positive predictions, resulting in a more favorable balance between sensitivity and precision. AUROC remained high for both approaches (0.999), while AUPRC increased from 0.949 to 0.962. These findings suggest that retrieval-augmented refinement enhances discriminative performance while maintaining strong overall classification performance. To assess robustness, the abstract selection and evaluation procedure was repeated four additional times using independently curated sets of manually verified disease pairs. Consistent performance trends were observed across all repetitions, with detailed results provided in the Supplementary Table 2-5.

**Table 6.** Test Results for RAER Compared with the Zero-Shot Baseline.

| Model/Configuration | Accuracy | Precision | Recall | F1 | AUROC | AUPRC |
| --- | --- | --- | --- | --- | --- | --- |
| Zero-shot | 0.992 | 0.674 | **0.967** | 0.795 | 0.999 | 0.949 |
| k8_τ0.70_β9_α0.75 | **0.994** | **0.778** | 0.933 | **0.848** | 0.999 | **0.962** |
| k8_τ0.70_β12_α0.75 | 0.998 | 0.778 | 0.933 | 0.848 | 0.999 | 0.962 |
| k8_τ0.70_β15_α0.75 | 0.998 | 0.778 | 0.933 | 0.848 | 0.999 | 0.961 |
| k10_τ0.70_β9_α0.75 | 0.998 | 0.778 | 0.933 | 0.848 | 0.999 | 0.962 |
| k10_τ0.70_β12_α0.75 | 0.998 | 0.778 | 0.933 | 0.848 | 0.999 | 0.962 |
| k10_τ0.70_β15_α0.75 | 0.998 | 0.778 | 0.933 | 0.848 | 0.999 | 0.961 |

Overall, RAER consistently outperformed zero-shot embeddings across both parameter optimization and independent testing. The framework demonstrated stable improvements in precision, F1-score, and AUPRC while maintaining high accuracy and AUROC, supporting retrieval-augmented embedding refinement as an effective approach for rare disease text matching without task-specific fine-tuning.

## Discussion

In this study, we developed RDEmbed, a sentence-embedding framework for rare disease phenotyping and phenotypic similarity analysis and systematically compared embedding-based similarity measures with traditional HPO-driven semantic similarity scores. Our results demonstrate that sentence-level embeddings, particularly highest-match measures (cosine_top5, cosine_topk), outperformed ontology-based approaches in identifying phenotypically similar case reports across both simulated and real datasets.

This finding highlights the potential of embedding-based approaches to enhance rare disease diagnosis, patient matchmaking, and discovery of novel disease associations. Building on these findings, the proposed RAER framework further extends embedding-based phenotypic representation through retrieval-augmented contextual refinement, yielding improved discriminative performance over zero-shot embeddings on the independent test set. This indicates a trade-off toward higher specificity in rare disease sentence matching while preserving strong overall classification performance. HPO-based semantic similarity measures such as OWLSim and simGIC have proven useful for systematic comparison of patient phenotypes and gene prioritization [16], [18]. However, these methods are limited by the accuracy of HPO term extraction from unstructured text, incompleteness of annotations, and the inherent constraints of information content–based metrics in capturing nuanced semantic context [19], [21]. Our results corroborate previous observations that ontology-based approaches, while robust for structured datasets, can miss subtle but clinically important phenotypic relationships, reducing sensitivity in rare disease identification[42].

Transformer-based sentence embeddings offer several advantages over ontology-driven methods. By representing clinical narratives in a continuous semantic vector space, embeddings capture contextual meaning beyond pre-defined ontology terms [25], [26]. This allows identification of phenotypically similar cases even when explicit HPO annotations are incomplete or absent. Our findings align with emerging literature on embedding-based biomedical text mining. Domain-adapted transformer models such as BioBERT and ClinicalBERT have shown state-of-the-art performance in tasks ranging from named entity recognition to phenotypic similarity analysis [23], [24]. Recent work applying sentence embeddings to clinical narratives has highlighted their potential for rare disease phenotyping and cohort discovery [26]. By directly comparing embedding-based similarity with HPO-driven methods on both simulated and real case report datasets, our study provides quantitative evidence that embeddings offer a more sensitive and generalizable approach for phenotypic comparison.

Nevertheless, some limitations should be considered while interpreting these results in conjunction with the existing HPO-based approaches. First, although sentence embeddings capture rich semantic context, their performance may vary depending on the quality of input narratives and model pre-training. Very short or poorly detailed abstracts (or patient summaries) may yield less informative embeddings, potentially affecting similarity calculations. Second, our real-world evaluation relied on PubMed abstracts, which may not fully capture the depth of electronic health record data. We do acknowledge that our testing datasets comprise highly controlled datasets that the transformer-based approaches can substantially thrive due to strong contextual cues that they can extract. Furthermore, our approach used an automated HPO extraction, which may not fully capitulate. Finally, while embeddings improve precision, HPO-based approaches retain advantages in interpretability and linkage to curated ontological knowledge, suggesting that hybrid strategies combining both methods may offer complementary benefits for rare disease research.

These limitations highlight important directions for future research. Building on the promising performance of RDEmbed in case report abstracts, a natural extension is its application to electronic health record (EHR) datasets. These narratives often contain rich, longitudinal phenotypic information, including comorbidities, laboratory findings, procedures, and treatment responses, which are not fully captured in PubMed abstracts. Applying RDEmbed to EHR datasets could enable large-scale phenotypic similarity analyses, rare disease cohort identification, and longitudinal phenotyping, offering direct clinical utility for precision medicine. Challenges include handling heterogeneous formatting, varying levels of detail, and potential data sparsity, but transformer-based embeddings are well-suited to capture contextual semantics across diverse narrative structures. Future work may also explore hybrid approaches that integrate embeddings with HPO term extraction to combine interpretability with contextual richness, improving both diagnostic support and rare disease discovery in real-world clinical settings.

## Acknowledgements

AH, XW, JW, HL, and ZZ conceived and designed the study. XW and JW designed and performed the experiments, collected and visualized the results. All authors wrote and approved the manuscript. AH and XW were supported by funding from National Institutes of Health.

## Funding Declaration

AH and XW were supported by funding from National Institutes of Health grant number R01HG012604 and discretionary funds from The University of Texas Health Science Center Houston.

## Ethics Declaration

Not applicable

## Methods

### Datasets and Resources

A simulated dataset of PubMed-style case report abstracts was generated for 100 rare diseases selected from the Orphanet database using the OpenAI GPT-4o model. For each disease, two distinct abstracts were simulated, yielding a total of 200 case reports. Abstracts are organized in consecutive pairs (e.g., 1–2, 3–4, …, 199–200), with each pair representing two independent case descriptions of the same rare disease. Each abstract comprised a structured *Title* and *Abstract* section, with deliberate variation in title format, patient demographics, clinical presentations, diagnostic findings, therapeutic interventions, and outcomes to ensure diversity. This dataset provides a standardized resource for benchmarking phenotyping approaches, facilitating systematic evaluation of the RDEmbed method relative to traditional HPO-based phenotyping techniques.

To assess RDEmbed performance on real-world data, PubMed was systematically queried using structured URL-based search requests with predefined keywords “unidentified case study”. Searches were restricted to case reports published within the past five years and with available abstracts, yielding 2,080 records. Abstracts containing fewer than 180 words were excluded to ensure adequate clinical detail, resulting in a subset of 496 abstracts. From this subset, 200 abstracts were randomly selected for verification, enabling comparative assessment of RDEmbed-based phenotyping against conventional HPO-based methods in accurately matching the associated rare diseases. Additionally, we incorporated a case-study corpus consisting of 156 clinical abstracts representing six rare diseases, curated by our team, to assess disease-specific clustering performance and to further validate the utility of RDEmbed for phenotypic similarity analysis. The dataset comprises Cystinuria (7 abstracts), Dent disease (11 abstracts), Primary Hyperoxaluria (PH; 13 abstracts), Idiopathic Pulmonary Fibrosis (IPF; 19 abstracts), Mastocytosis (47 abstracts), and Hypereosinophilic Syndrome (HES; 59 abstracts), totaling 156 entries.

To evaluate the proposed retrieval-augmented embedding refinement (RAER) framework, A benchmark dataset of 30 rare diseases was constructed using PubMed abstracts published between 2022 and 2026. This period was chosen to evaluate knowledge acquired after the training of the all-MiniLM-L6-v2 embedding model, which was pretrained primarily on pre-2021 literature. For each disease, two abstracts were used for parameter optimization and two for independent testing, with all positive disease pairs manually verified. An additional 30 disease-related abstracts per disease were incorporated into a vector database to support retrieval-augmented embedding refinement. To assess the robustness of the proposed framework, the abstract selection and evaluation process was repeated four additional times using independent sets of manually verified disease pairs.

### HPO-based phenotyping

In this study, we employed PhenoBERT [32], a deep learning approach for automated extraction of HPO terms from unstructured text. PhenoBERT has demonstrated superior performance compared with other deep learning methods, such as NeuralCR [43] and PhenoTagger [44], particularly in more challenging prediction tasks, and offers greater computational efficiency.

Phenotypic similarity scores were calculated using the PhenomeCentral package [18], which applies HPO-based semantic similarity measures. Specifically, the OWLSim score and simGIC score was employed to quantify pairwise similarity between patients. OWLSim quantifies semantic similarity between two sets of HPO terms by performing a pairwise comparison of all terms and aggregating the highest-scoring matches. Similarity between individual terms is computed using either the Jaccard index or information content (IC), and the overall score reflects the average of the best-matching pairs across the two sets [15]. OWLSim has been integrated with tools such as Exomiser [16], [18] and PhenoDigm [17] for large-scale phenotype analyses. Whereas simGIC quantifies phenotypic similarity as the ratio of the summed IC of shared HPO terms to the total IC of all HPO terms in the two sets, it is formally defined as:

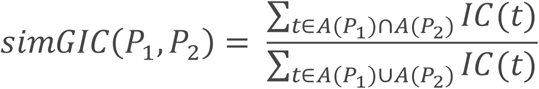

where *A*(*P*_1_) and *A*(*P*_2_) denote the sets of all ancestor HPO terms (including the original terms) for patients *P*_1_and *P*_2_, and *IC*(*t*) represents the information content of term t [14]. SimGIC score demonstrated superior performance, consistently ranking the correct match highly across various methods for calculating information content (IC) [18].

### Embedding-based phenotyping

For our RDEmbed, the embedding-based phenotyping, each abstract was first segmented into individual sentences and encoded into 384-dimensional vectors using the pre-trained Sentence-Transformer model all-MiniLM-L6-v2 (https://huggingface.co/sentence-transformers/all-MiniLM-L6-v2). Let 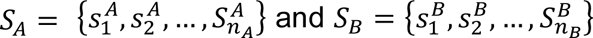 denote the sets of sentence embeddings for patients *A* and *B*, respectively, and let cos(*s_i_*, *s_j_*) represent the cosine similarity between two sentence embeddings.

The average cosine similarity (cosine_avg) across all sentence pairs was calculated as:

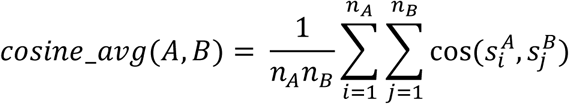

To highlight the most informative semantic matches between abstracts, the top-5 and top-10 similarities were computed by first collecting all pairwise sentence similarities into a single list and sorting them in descending order. Let *sorted*_*maxSim* denote this sorted list of pairwise similarities, so that the first element corresponds to the maximum similarity between any sentence pair:

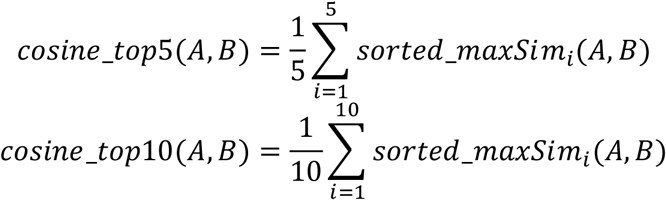

To account for differences in abstract length and sentence number, two complementary approaches were implemented:

1. Bidirectional maximum similarity (cosine_bi): For each sentence in abstract *A*, the maximum similarity with any sentence in *B* was computed, and similarly for each sentence in *B* with sentences in *A*. The bidirectional similarity is then obtained by averaging both sets:

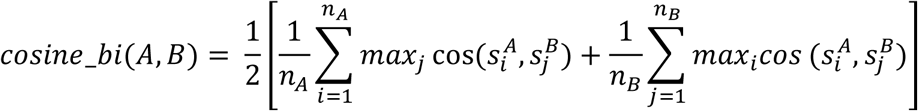

1. Dynamic top-k similarity (cosine_topk): The top *k* values from the sorted list of all pairwise sentence similarities (*sorted*_*maxSim*) are averaged. The number of matches *k* is set to 75% of the shorter abstract’s length, bounded between 3 and 15 sentences.

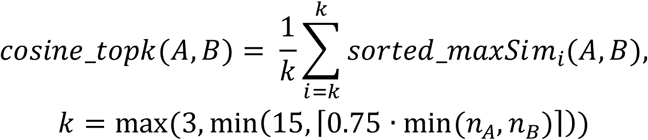

These complementary metrics provide a robust framework for quantitative comparison of semantically similar clinical descriptions, facilitating phenotype-based patient matching and downstream analyses.

### Retrieval-Augmented Embedding Refinement

To incorporate rare disease specific knowledge into the embedding representation, we developed a retrieval-augmented embedding refinement (RAER) framework. Given an input sentence, the zero-shot embedding vector (*e*_0_) was first generated using the pretrained all-MiniLM-L6-v2 sentence embedding model. The embedding was then used to query a vector database containing the disease-related sentences and the top-*k* nearest neighbors were retrieved based on the cosine similarity. The final retrieval-augmented embedding was computed as:

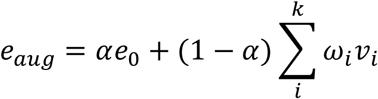

where *v_i_* denotes the embedding of the *i*-th retrieved sentence and *α* controls the relative contribution of the original embedding and the retrieved contextual information. The similarity weights *ω_i_* were computed using a temperature-scaled softmax function:

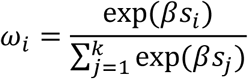

where *s_i_* is the cosine similarity between the query embedding and the *i*-th retrieved embedding, and *β* is a temperature parameter controlling the concentration of the weight distribution.

To ensure that only semantically relevant retrieved sentences contribute to the final representation, a similarity threshold *τ* is applied to filter out neighbors with low cosine similarity to the query. Specifically, retrieved sentences with similarity below *τ* are excluded from downstream aggregation, thereby constraining the retrieval context to high-confidence neighbors. The final retrieval context embedding is computed using a similarity-weighted aggregation of the remaining neighbors.

Hyperparameters, including the number of retrieved neighbors *k* ∈ {5, 8, 10}, similarity threshold *τ* ∈ {0.65, 0.70, 0.75}, temperature parameter *β* ∈ {9, 12, 15}, and aggregation weight *α* ∈ {0.75, 0.80, 0.85}, were tuned on a dedicated parameter optimization dataset. A greedy search strategy was employed for hyperparameter selection, sequentially optimizing each parameter to identify the best-performing configuration. The resulting optimal parameter set was then applied to the held-out testing dataset for final evaluation.

### Evaluation and Performance Metrics

The performance of rare disease identification was evaluated using both HPO-based semantic similarity scores and embedding-based cosine similarity scores. Sentence embeddings were visualized with t-SNE to examine abstract clustering, with abstracts exhibiting higher cosine similarity clustering together. For comparison, all unique HPO terms across abstracts were converted into a binary presence/absence matrix (one-hot encoding) for downstream analysis. This matrix was also subjected to t-SNE, projecting high-dimensional phenotypic profiles into two dimensions while preserving local structure, enabling visualization of clustering patterns among abstracts with similar phenotypes. Our performance metrics include the proportion of cases in which the true match ranked first or within the top five matches, as well as ROC and PRC curves, accuracy, precision, recall, and F1-score, calculated for the simulated abstracts. To validate the methods on real abstracts, thresholds were determined from the ROC curves for the top-performing scores of each approach, simGIC for HPO-based phenotyping and cosine_top5 similarity for embedding-based phenotyping, and applied to identify matching rare diseases, with results compared across methods.

### Case Study

To further validate our embedding-based phenotyping approach, we evaluated its performance against HPO-based methods using a curated dataset comprising 156 clinical abstracts across six rare diseases. We assessed model performance through t - SNE visualizations and quantitative metrics including receiver operating characteristic (ROC) and precision–recall (PRC) curves, as well as accuracy, precision, recall, and F1-score.

## Notes

### Competing Interest Statement

The authors have declared no competing interest.

