## Supplemental data for "RDEmbed: A Sentence-Embedding Framework for Rare Disease Phenotype Representation and Phenotypic Similarity Analysis"

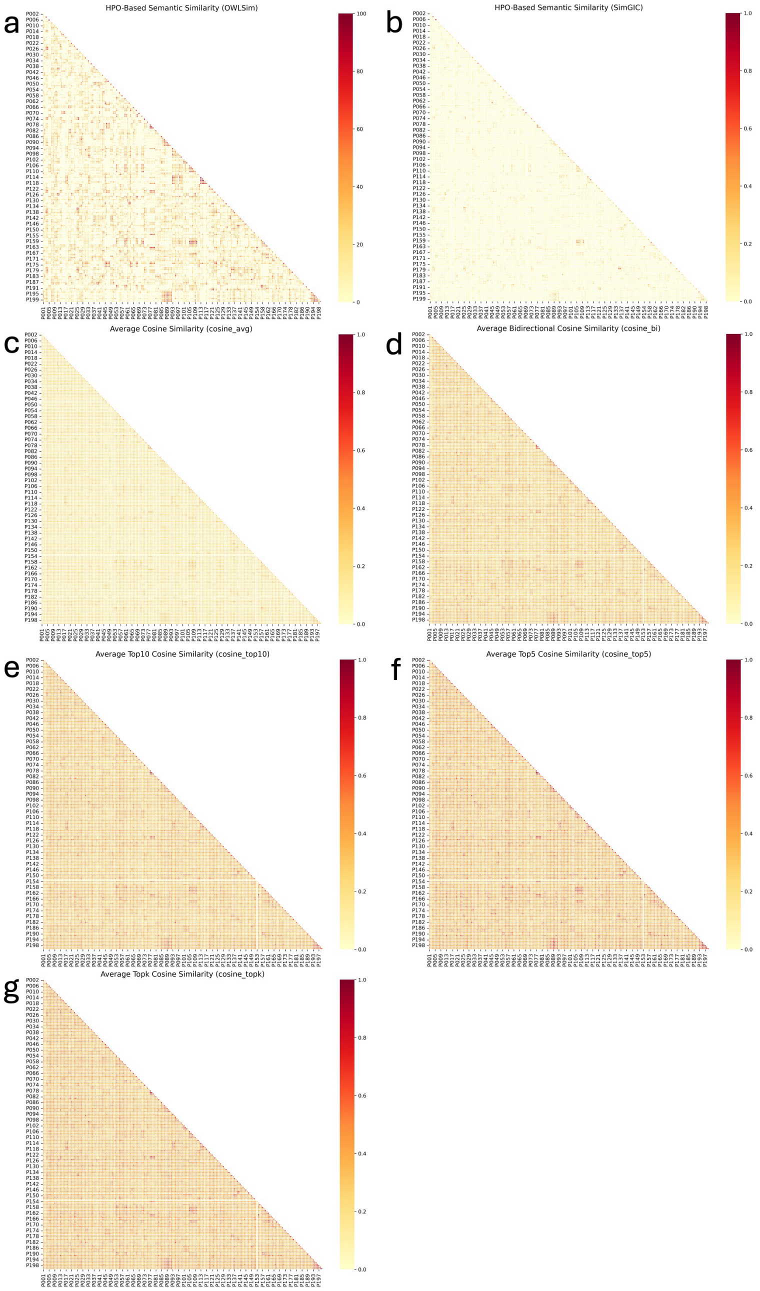


**Fig. S1. Heat map of seven phenotyping similarity score matrices for 100 pairs of simulated abstracts.** Abstracts are arranged in consecutive pairs (1–2, 3–4, …, 199–200), with each pair representing two independent case descriptions of the same rare disease. Higher similarity scores are evident along the diagonal, reflecting strong within-pair concordance. **(a)** HPO-based semantic similarity (OWLSim); **(b)** HPO-based semantic similarity (SimGIC); **(c)** Average cosine similarity (cosine_avg) computed with RDEmbed; **(d)** Bidirectional average cosine similarity (cosine_bi) computed with RDEmbed; **(e)** Top10 average cosine similarity (cosine_top10) computed with RDEmbed; **(f)** Top5 average cosine similarity (cosine_top5) computed with RDEmbed; and **(g)** Dynamic topk average cosine similarity (cosine_topk) computed with RDEmbed.

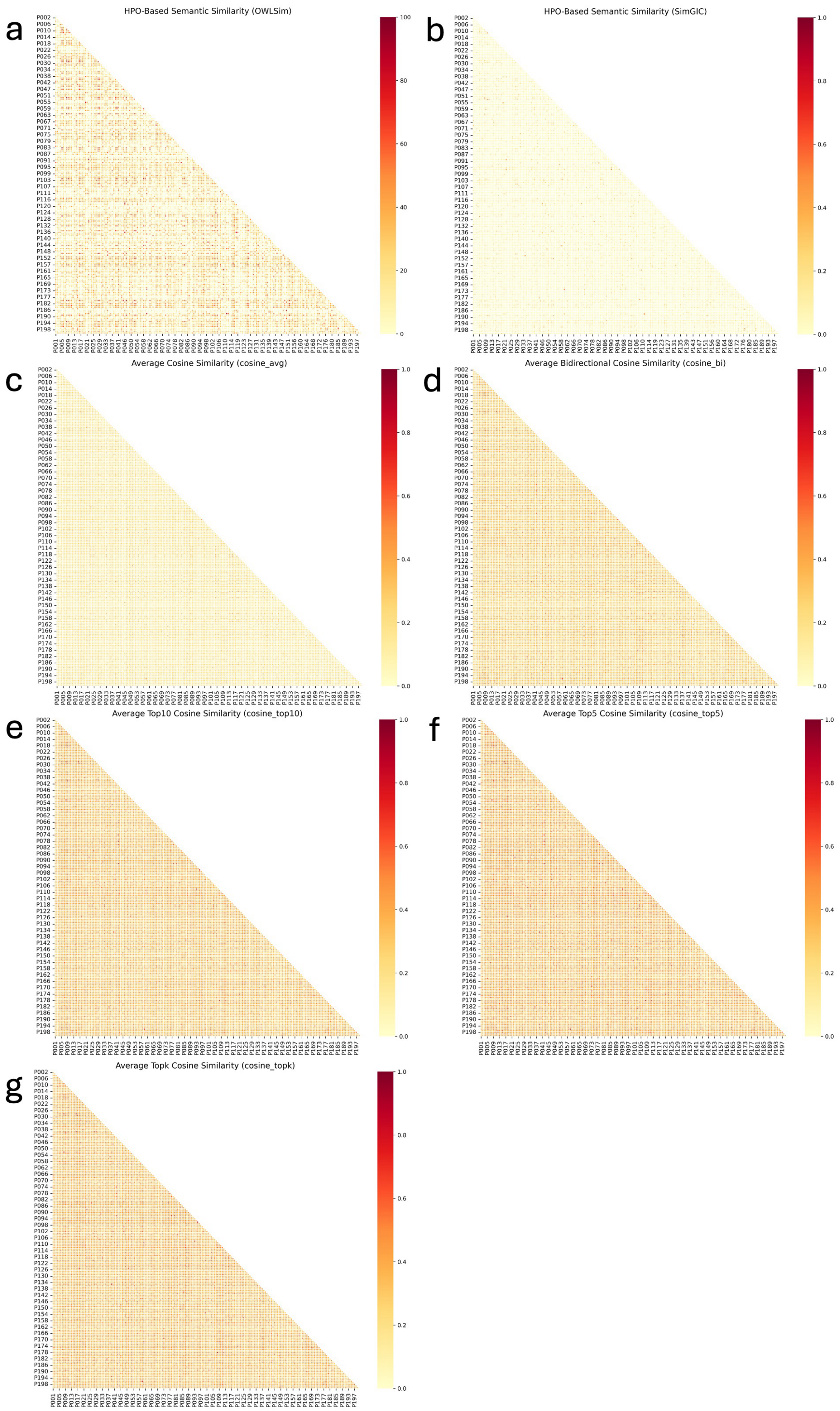


**Fig. S2. Heat map of seven phenotyping similarity score matrices for 200 real PubMed abstracts.** **(a)** HPO-based semantic similarity (OWLSim); **(b)** HPO-based semantic similarity (SimGIC); **(c)** Average cosine similarity (cosine_avg) computed with RDEmbed; **(d)** Bidirectional average cosine similarity (cosine_bi) computed with RDEmbed; **(e)** Top10 average cosine similarity (cosine_top10) computed with RDEmbed; **(f)** Top5 average cosine similarity (cosine_top5) computed with RDEmbed; and **(g)** Dynamic topk average cosine similarity (cosine_topk) computed with RDEmbed.

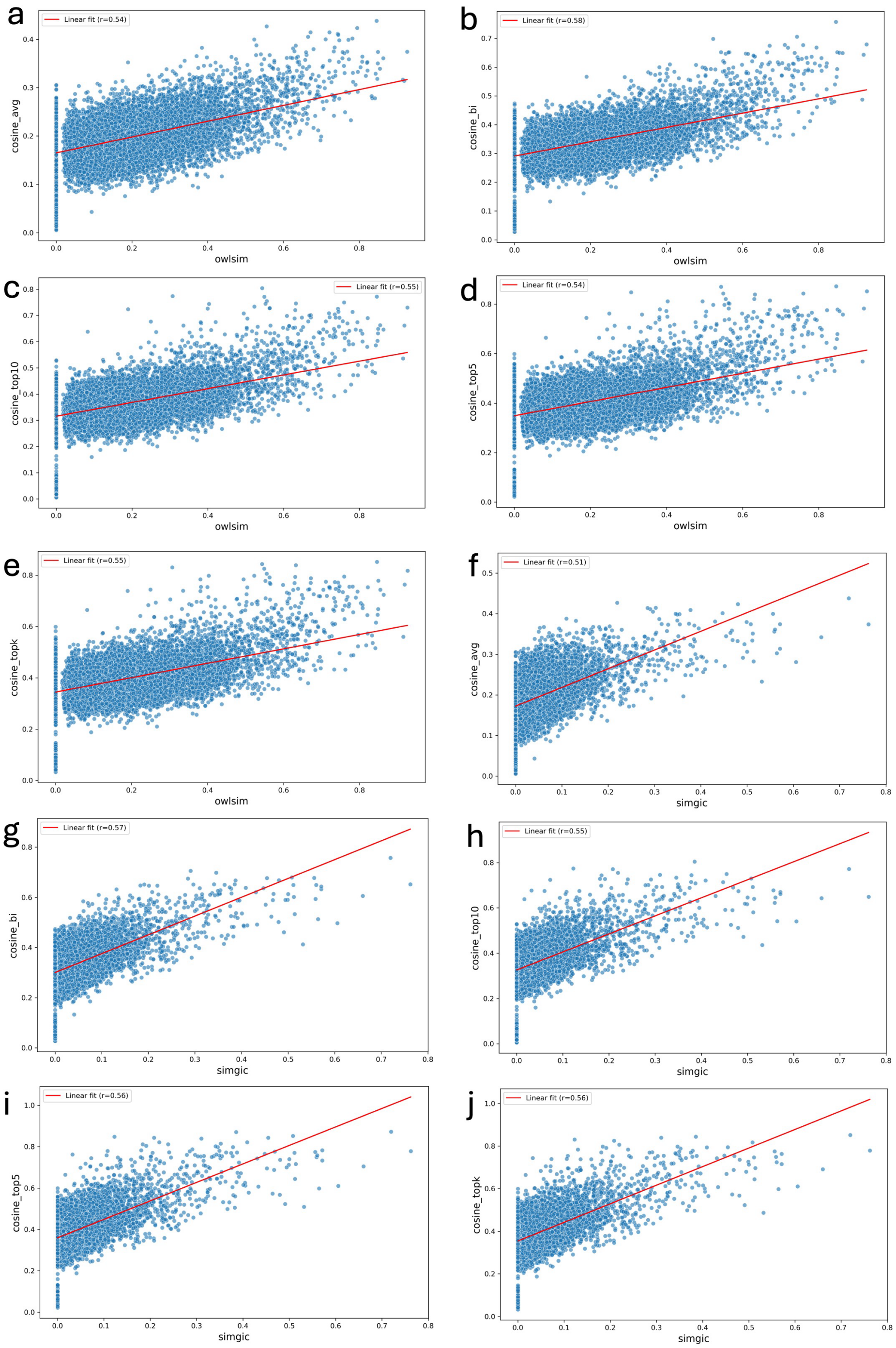


**Fig. S3. Correlation between HPO-based semantic similarity scores and RDEmbed cosine similarity scores in 100 pairs of simulated abstracts.** Scatterplots illustrate pairwise correlations between the HPO-based measures OWLSim and SimGIC and five RDEmbed cosine similarity metrics (cosine_avg, cosine_bi, cosine_top10, cosine_top5, and cosine_topk). Overall correlations were moderate (r = 0.51–0.58), indicating that the two approaches capture overlapping phenotypic information. **(a)** OWLSim vs. cosine_avg; **(b)** OWLSim vs. cosine_bi; **(c)** OWLSim vs. cosine_top10; **(d)** OWLSim vs. cosine_top5; **(e)** OWLSim vs. cosine_topk; **(f)** SimGIC vs. cosine_avg; **(g)** SimGIC vs. cosine_bi; **(h)** SimGIC vs. cosine_top10; **(i)** SimGIC vs. cosine_top5; and **(j)** SimGIC vs. cosine_topk.

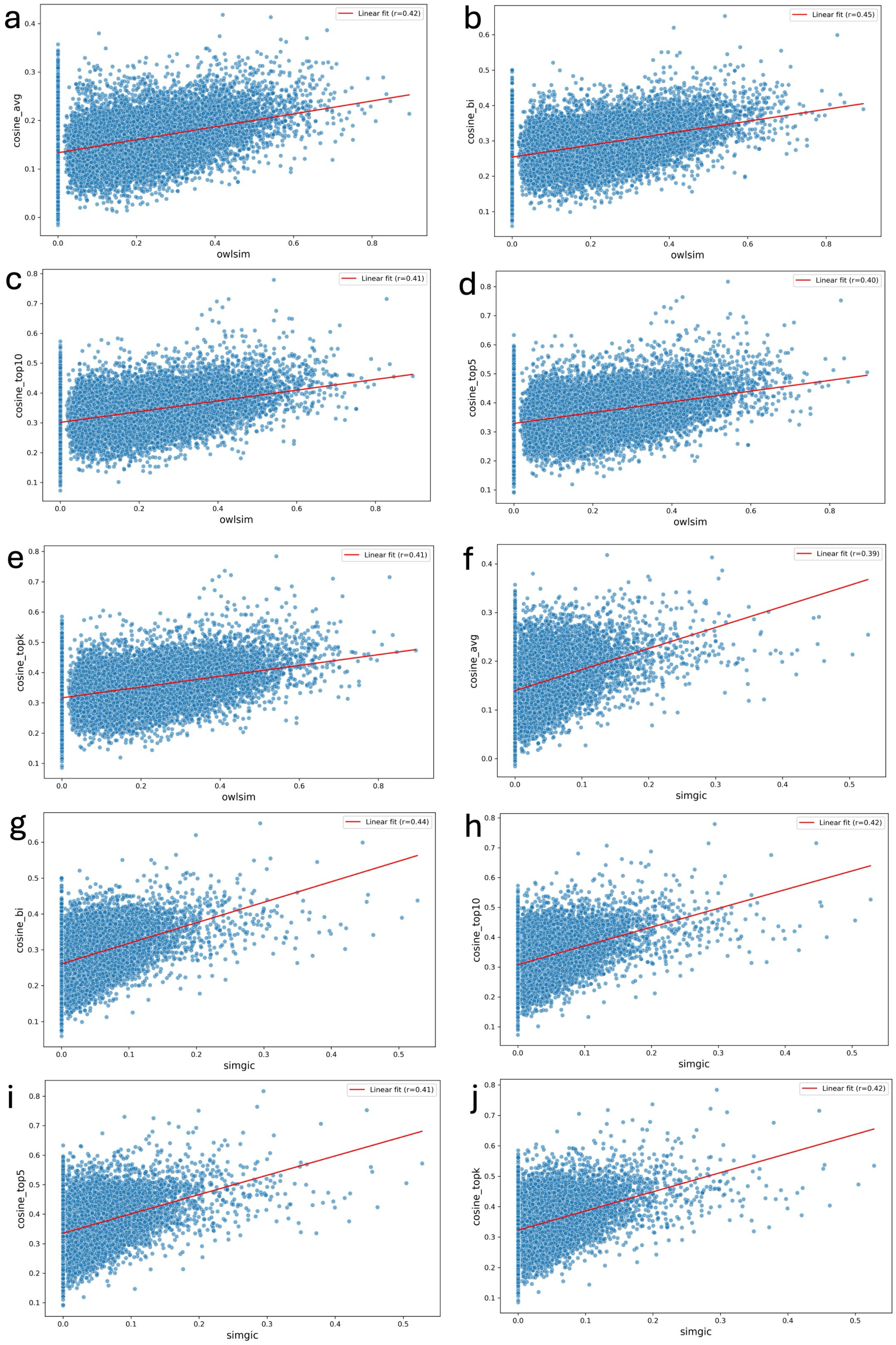


**Fig. S4. Correlation between HPO-based semantic similarity scores and RDEmbed cosine similarity scores in 200 real PubMed abstracts.** Scatterplots illustrate pairwise correlations between the HPO-based measures OWLSim and SimGIC and five RDEmbed cosine similarity metrics (cosine_avg, cosine_bi, cosine_top10, cosine_top5, and cosine_topk). Overall correlations were moderate (r = 0.39–0.45), indicating that the two approaches capture overlapping phenotypic information. **(a)** OWLSim vs. cosine_avg; **(b)** OWLSim vs. cosine_bi; **(c)** OWLSim vs. cosine_top10; **(d)** OWLSim vs. cosine_top5; **(e)** OWLSim vs. cosine_topk; **(f)** SimGIC vs. cosine_avg; **(g)** SimGIC vs. cosine_bi; **(h)** SimGIC vs. cosine_top10; **(i)** SimGIC vs. cosine_top5; and **(j)** SimGIC vs. cosine_topk.

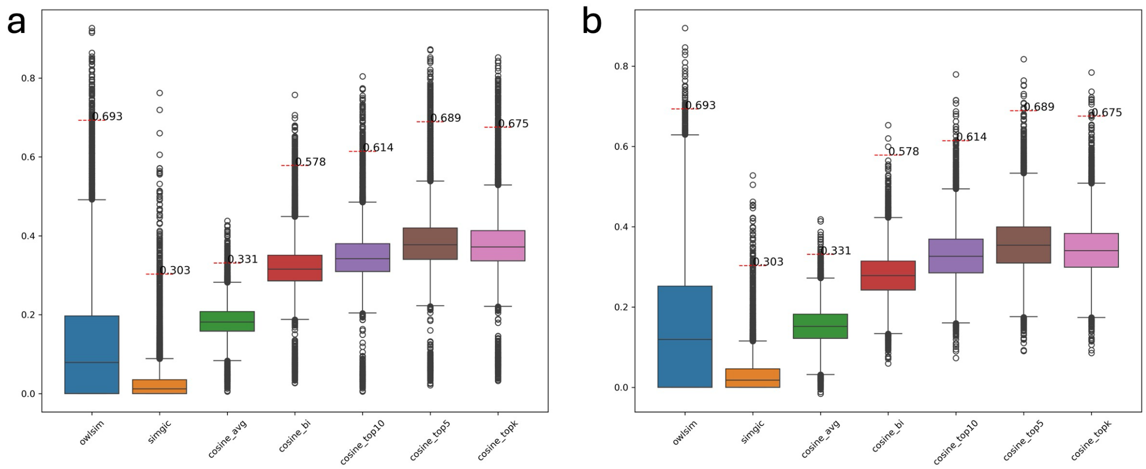


**Fig. S5. Box plots of seven phenotyping similarity scores with optimal decision thresholds.** Distributions of seven similarity measures, owlsim, simgic, cosine_avg, cosine_bi, cosine_top10, cosine_top5, and cosine_topk, are shown as box plots. Horizontal dashed lines indicate the optimal decision thresholds derived from the corresponding ROC curves. **(a)** 100 pairs of simulated abstracts; **(b)** 200 real PubMed abstracts.

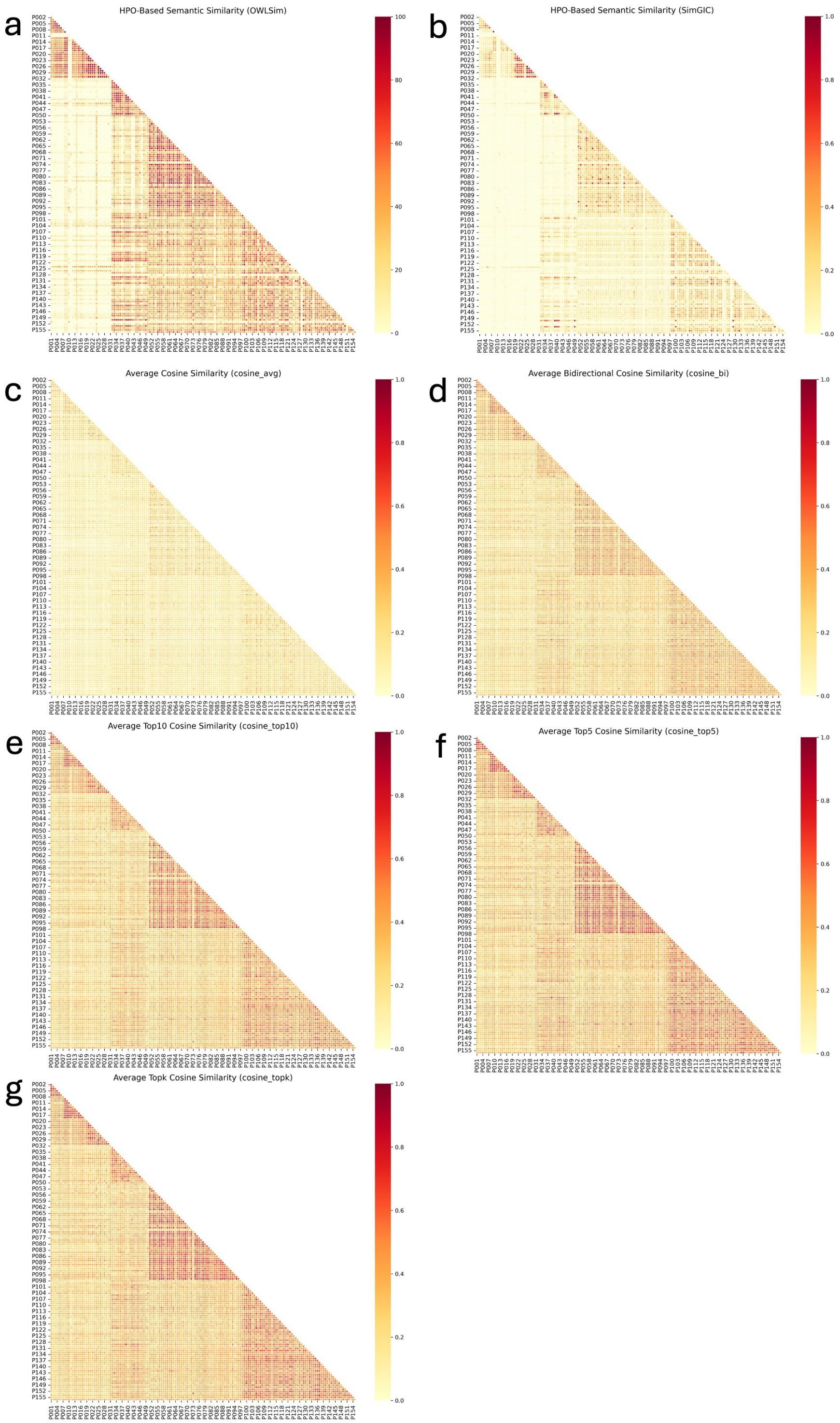


**Fig. S6. Heat map of seven phenotyping similarity score matrices for six groups of rare disease abstracts. (a)** HPO-based semantic similarity (OWLSim); **(b)** HPO-based semantic similarity (SimGIC); **(c)** Average cosine similarity (cosine_avg) computed with RDEmbed; **(d)** Bidirectional average cosine similarity (cosine_bi) computed with RDEmbed; **(e)** Top10 average cosine similarity (cosine_top10) computed with RDEmbed; **(f)** Top5 average cosine similarity (cosine_top5) computed with RDEmbed; and **(g)** Dynamic topk average cosine similarity (cosine_topk) computed with RDEmbed.

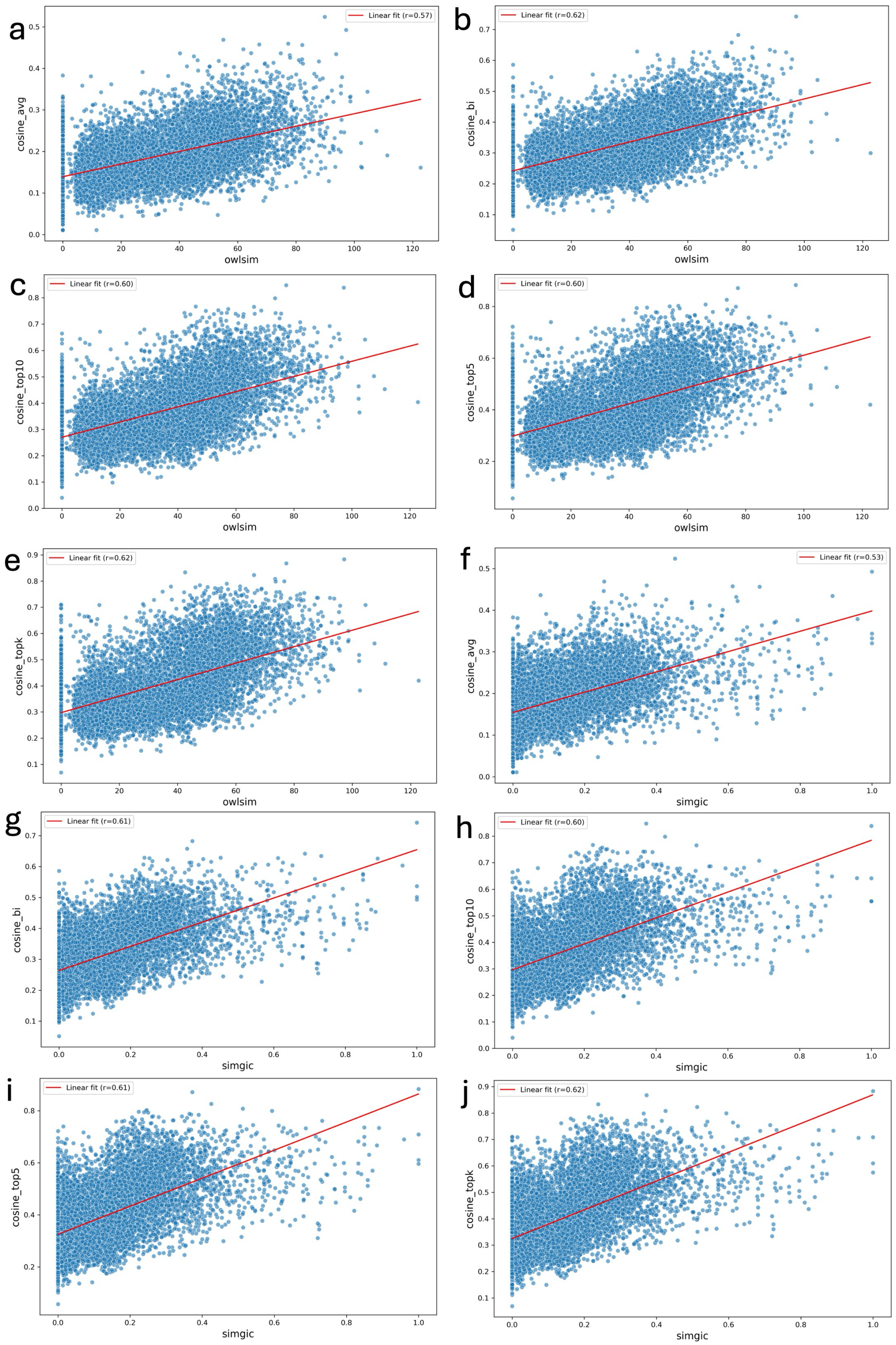


**Fig. S7. Correlation between HPO-based semantic similarity scores and RDEmbed cosine similarity scores for six groups of rare disease abstracts.** Scatterplots illustrate pairwise correlations between the HPO-based measures OWLSim and SimGIC and five RDEmbed cosine similarity metrics (cosine_avg, cosine_bi, cosine_top10, cosine_top5, and cosine_topk). Overall correlations were moderate (r = 0.53–0.62), indicating that the two approaches capture overlapping phenotypic information. **(a)** OWLSim vs. cosine_avg; **(b)** OWLSim vs. cosine_bi; **(c)** OWLSim vs. cosine_top10; **(d)** OWLSim vs. cosine_top5; **(e)** OWLSim vs. cosine_topk; **(f)** SimGIC vs. cosine_avg; **(g)** SimGIC vs. cosine_bi; **(h)** SimGIC vs. cosine_top10; **(i)** SimGIC vs. cosine_top5; and **(j)** SimGIC vs. cosine_topk.

**Table S2a.** RAER Parameter Tuning Results Compared with the Zero-Shot Baseline (repeat dataset 1)

| Model/Configuration | Threshold | Accuracy | Precision | Recall | F1-score | AUROC | AUPRC |
|---|---|---|---|---|---|---|---|
| Zero-shot | 0.470 | 0.968 | 0.349 | 1.0 | 0.517 | 0.998 | 0.945 |
| k8_τ0.65_β9_α0.75 | 0.542 | **0.989** | 0.600 | 1.0 | 0.750 | 0.999 | **0.960** |
| k8_τ0.65_β12_α0.75 | 0.542 | 0.989 | **0.612** | 1.0 | **0.759** | 0.999 | 0.960 |
| k8_τ0.65_β15_α0.75 | 0.542 | 0.989 | 0.600 | 1.0 | 0.750 | 0.999 | 0.960 |
| k10_τ0.65_β15_α0.75 | 0.542 | 0.989 | 0.600 | 1.0 | 0.750 | 0.999 | 0.960 |

**Table S2b.** RAER Test Results Compared with the Zero-Shot Baseline (repeat dataset 1)

| Model/Configuration | Accuracy | Precision | Recall | F1-score | AUROC | AUPRC |
|---|---|---|---|---|---|---|
| Zero-shot | 0.993 | 0.725 | **0.967** | 0.829 | 0.998 | 0.943 |
| k8_τ0.65_β9_α0.75 | **0.994** | **0.771** | 0.900 | **0.831** | **0.998** | **0.953** |
| k8_τ0.65_β12_α0.75 | 0.994 | 0.771 | 0.900 | 0.831 | 0.998 | 0.953 |
| k8_τ0.65_β15_α0.75 | 0.994 | 0.771 | 0.900 | 0.831 | 0.998 | 0.952 |
| k10_τ0.65_β15_α0.75 | 0.994 | 0.771 | 0.900 | 0.831 | 0.998 | 0.953 |

**Table S3a.** RAER Parameter Tuning Results Compared with the Zero-Shot Baseline (repeat dataset 2)

| Model/Configuration | Threshold | Accuracy | Precision | Recall | F1-score | AUROC | AUPRC |
|---|---|---|---|---|---|---|---|
| Zero-shot | 0.475 | 0.985 | 0.526 | 1.0 | 0.690 | 0.999 | 0.972 |
| k5_τ0.65_β9_α0.75 | 0.554 | **0.998** | **0.909** | 1.0 | **0.952** | **1.0** | **0.984** |
| k5_τ0.65_β12_α0.75 | 0.551 | 0.998 | 0.909 | 1.0 | 0.952 | 1.0 | 0.984 |
| k8_τ0.65_β9_α0.75 | 0.554 | 0.998 | 0.909 | 1.0 | 0.952 | 1.0 | 0.984 |
| k8_τ0.65_β12_α0.75 | 0.552 | 0.998 | 0.909 | 1.0 | 0.952 | 1.0 | 0.984 |
| k10_τ0.65_β9_α0.75 | 0.555 | 0.998 | 0.909 | 1.0 | 0.952 | 1.0 | 0.984 |
| k10_τ0.65_β12_α0.75 | 0.552 | 0.998 | 0.909 | 1.0 | 0.952 | 1.0 | 0.984 |

**Table S3b.** RAER Test Results Compared with the Zero-Shot Baseline (repeat dataset 2)

| Model/Configuration | Accuracy | Precision | Recall | F1-score | AUROC | AUPRC |
|---|---|---|---|---|---|---|
| Zero-shot | 0.988 | 0.577 | **1.0** | 0.732 | 0.999 | 0.992 |
| k5_τ0.65_β9_α0.75 | 0.998 | **0.935** | 0.967 | **0.951** | **0.999** | **0.995** |
| k5_τ0.65_β12_α0.75 | 0.998 | 0.935 | 0.967 | 0.951 | 0.999 | 0.995 |
| k8_τ0.65_β9_α0.75 | 0.998 | 0.935 | 0.967 | 0.951 | 0.999 | 0.995 |
| k8_τ0.65_β12_α0.75 | 0.998 | 0.935 | 0.967 | 0.951 | 0.999 | 0.994 |
| k10_τ0.65_β9_α0.75 | 0.998 | 0.935 | 0.967 | 0.951 | 0.999 | 0.995 |
| k10_τ0.65_β12_α0.75 | 0.998 | 0.935 | 0.967 | 0.951 | 0.999 | 0.995 |

**Table S4a.** RAER Parameter Tuning Results Compared with the Zero-Shot Baseline (repeat dataset 3)

| Model/Configuration | Threshold | Accuracy | Precision | Recall | F1-score | AUROC | AUPRC |
|---|---|---|---|---|---|---|---|
| Zero-shot | 0.477 | 0.977 | 0.418 | 0.933 | 0.577 | 0.986 | 0.855 |
| k5_τ0.75_β9_α0.75 | 0.498 | **0.986** | **0.560** | 0.933 | **0.700** | 0.989 | 0.873 |
| k5_τ0.75_β12_α0.75 | 0.498 | 0.986 | 0.560 | 0.933 | 0.700 | 0.989 | 0.873 |
| k5_τ0.75_β15_α0.75 | 0.497 | 0.986 | 0.560 | 0.933 | 0.700 | 0.989 | 0.872 |
| k8_τ0.65_β12_α0.75 | 0.492 | 0.965 | 0.318 | 0.933 | 0.475 | 0.990 | 0.875 |
| k8_τ0.75_β9_α0.75 | 0.498 | 0.986 | 0.560 | 0.933 | 0.700 | 0.989 | 0.874 |
| k8_τ0.75_β12_α0.75 | 0.498 | 0.986 | 0.560 | 0.933 | 0.700 | 0.989 | 0.873 |
| k8_τ0.75_β15_α0.75 | 0.497 | 0.986 | 0.560 | 0.933 | 0.700 | 0.989 | 0.873 |
| k10_τ0.65_β9_α0.75 | 0.493 | 0.966 | 0.322 | 0.933 | 0.479 | **0.990** | **0.876** |
| k10_τ0.65_β12_α0.75 | 0.494 | 0.966 | 0.326 | 0.933 | 0.483 | 0.990 | 0.875 |
| k10_τ0.75_β9_α0.75 | 0.499 | 0.986 | 0.560 | 0.933 | 0.700 | 0.989 | 0.875 |
| k10_τ0.75_β12_α0.75 | 0.498 | 0.986 | 0.560 | 0.933 | 0.700 | 0.989 | 0.875 |
| k10_τ0.75_β15_α0.75 | 0.498 | 0.986 | 0.560 | 0.933 | 0.700 | 0.989 | 0.874 |

**Table S4b.** RAER Test Results Compared with the Zero-Shot Baseline (repeat dataset 3)

| Model/Configuration | Accuracy | Precision | Recall | F1-score | AUROC | AUPRC |
|---|---|---|---|---|---|---|
| Zero-shot | 0.983 | 0.500 | 0.967 | 0.659 | 0.998 | 0.950 |
| k5_τ0.75_β9_α0.75 | **0.986** | **0.549** | 0.933 | **0.691** | 0.998 | 0.949 |
| k5_τ0.75_β12_α0.75 | 0.986 | 0.549 | 0.933 | 0.691 | 0.998 | 0.949 |
| k5_τ0.75_β15_α0.75 | 0.986 | 0.549 | 0.933 | 0.691 | 0.998 | 0.949 |
| k8_τ0.65_β12_α0.75 | 0.978 | 0.435 | **1.0** | 0.606 | **0.999** | **0.954** |
| k8_τ0.75_β9_α0.75 | 0.986 | 0.549 | 0.933 | 0.691 | 0.998 | 0.949 |
| k8_τ0.75_β12_α0.75 | 0.986 | 0.549 | 0.933 | 0.691 | 0.998 | 0.949 |
| k8_τ0.75_β15_α0.75 | 0.986 | 0.549 | 0.933 | 0.691 | 0.998 | 0.949 |
| k10_τ0.65_β9_α0.75 | 0.978 | 0.435 | 1.0 | 0.606 | 0.999 | 0.954 |
| k10_τ0.65_β12_α0.75 | 0.978 | 0.435 | 1.0 | 0.606 | 0.999 | 0.954 |
| k10_τ0.75_β9_α0.75 | 0.986 | 0.549 | 0.933 | 0.691 | 0.998 | 0.949 |
| k10_τ0.75_β12_α0.75 | 0.986 | 0.549 | 0.933 | 0.691 | 0.998 | 0.949 |
| k10_τ0.75_β15_α0.75 | 0.986 | 0.549 | 0.933 | 0.691 | 0.998 | 0.949 |

**Table S5a.** RAER Parameter Tuning Results Compared with the Zero-Shot Baseline (repeat dataset 4)

| Model/Configuration | Threshold | Accuracy | Precision | Recall | F1-score | AUROC | AUPRC |
|---|---|---|---|---|---|---|---|
| Zero-shot | 0.426 | 0.928 | 0.191 | 1.0 | 0.321 | 0.994 | 0.885 |
| k8_τ0.65_β9_α0.75 | 0.462 | 0.949 | 0.250 | 1.0 | 0.400 | 0.995 | 0.898 |
| k8_τ0.70_β9_α0.75 | 0.449 | **0.950** | **0.252** | 1.0 | **0.403** | **0.995** | 0.901 |
| k10_τ0.70_β9_α0.75 | 0.449 | 0.949 | 0.250 | 1.0 | 0.400 | 0.995 | **0.902** |
| k10_τ0.70_β12_α0.75 | 0.449 | 0.949 | 0.248 | 1.0 | 0.397 | 0.995 | 0.901 |
| k10_τ0.70_β15_α0.75 | 0.449 | 0.947 | 0.244 | 1.0 | 0.392 | 0.995 | 0.900 |

**Table S5b.** RAER Test Results Compared with the Zero-Shot Baseline (repeat dataset 4)

| Model/Configuration | Accuracy | Precision | Recall | F1-score | AUROC | AUPRC |
|---|---|---|---|---|---|---|
| Zero-shot | 0.941 | 0.215 | 0.933 | 0.350 | 0.992 | 0.800 |
| k8_τ0.65_β9_α0.75 | **0.957** | **0.275** | 0.933 | **0.424** | **0.993** | **0.812** |
| k8_τ0.70_β9_α0.75 | 0.954 | 0.262 | 0.933 | 0.409 | 0.992 | 0.797 |
| k10_τ0.70_β9_α0.75 | 0.954 | 0.262 | 0.933 | 0.409 | 0.992 | 0.799 |
| k10_τ0.70_β12_α0.75 | 0.954 | 0.262 | 0.933 | 0.409 | 0.992 | 0.800 |
| k10_τ0.70_β15_α0.75 | 0.954 | 0.262 | 0.933 | 0.409 | 0.992 | 0.799 |
